# Evidence for Independent Processing of Shape by Vision and Touch

**DOI:** 10.1101/2022.02.01.478696

**Authors:** Ryan L. Miller, David L. Sheinberg

## Abstract

Although visual object recognition is well-studied and relatively well understood, much less is understood about how shapes are recognized by touch and how such haptic stimuli might be compared to visual shapes. One might expect that the processes of visual and haptic object recognition engage identical brain structures given the similarity of the problem and the advantages of avoiding redundant brain circuitry, but it has not yet been established the extent to which this is true. We recruited human participants to perform a one-back same-different visual and haptic shape comparison task both within (i.e., comparing two visual shapes or two haptic shapes) and across (i.e., comparing visual with haptic shapes) modalities. Participants saw or felt a shape and responded according to whether they thought the shape was the same or different from the previously felt or seen shape. We then used various shape metrics to predict performance based on the shape, orientation, and modality of the two stimuli which were being compared on each trial. We found that the fixed orientation of the shape stimuli was an important factor for predicting within-modal behavior, but the orientation of shapes compared across modality did not depend on knowing the presented orientation. We also found that the metrics which best predict shape comparison behavior heavily depended on the modality of the two shapes. We take these results as evidence that object recognition is not necessarily performed in a single, modality-agnostic region.

## Introduction

Object recognition is a core capacity afforded by our visual system and, accordingly, has long been of great interest to psychologists, neuroscientists, and philosophers (Cheselden, 1728; Ettlinger, 1956; Bülthoff et al., 1995; Riesenhuber & Poggio, 1999; Martin, 2007; Peissig & Tarr, 2007; Marr, 2010). The ability to recognize objects is not exclusively a visual faculty, however; we are also quite adept at recognizing objects solely by touch when, for example, searching for a coin in a pocket or purse. Although we know a great deal about the role that somatosensory pathways play in the perception of basic dimensions of touch, such as texture and vibration (Klatzky et al., 1985; Lederman & Klatzky, 1987; Sathian, 2016), very little is understood about how the brain ultimately integrates this information to serve advanced functions such as haptic object recognition.

From one perspective, both visual and haptic object recognition might be assumed to be processed by shared neural circuitry. After all, object recognition taps the same basic ability, regardless of the source modality, and it would seem economical not to have duplicate machinery. Indeed, this view has received support from groups studying the human visual extrastriate regions such as lateral occipital cortex (LOC) and inferotemporal cortex (IT) using imaging (Grill-Spector et al., 1998; Amedi et al., 2001; Pietrini et al., 2004; Prather et al., 2004; Lee Masson et al., 2016), though also see Snow et al. (2015).

Alternatively, based on what we know about peripheral sensory transduction pathways, it is clear that visual and somatosensory signals originate from fundamentally different end organs and are, at least initially, processed independently. So, from this perspective, recognition of objects using information from these senses may be contained within their own modality-specific circuits. Further complicating the issue, we know it is possible to compare shapes which are perceived haptically with shapes perceived visually, meaning there must be some way for the neural machinery processing these unisensory stimuli to communicate shape information. If visual and haptic shapes are processed in the same areas, this comparison may be relatively simple. If processed separately, comparisons may only be possible through intermediaries such as classical association or prefrontal areas responsible for higher level cognition. Researchers investigating anterior intraparietal (AIP) and dorsolateral prefrontal cortex have found evidence to support this, finding these areas to be especially active when comparing shapes across modalities (Murata et al., 2000; Grefkes et al., 2002; Ricciardi et al., 2006; Lacey et al., 2010; Helbig et al., 2012).

Here we attempt to disentangle these possibilities by determining the extent to which shapes from the two modalities are similarly represented. Figure 1 illustrates two alternative hypotheses which we are seeking to differentiate. Following basic feature extraction in visual and somatosensory unisensory areas, little is known about how haptic object recognition is completed and how much of the related neural machinery overlaps with what are considered visual processing areas. To the extent that there is a great deal of overlap (“early-convergence model”), we might expect that the same shape features determined to be crucial for visual object recognition would also be crucial for haptic object recognition. On the other hand, if the properties which are relevant for recognizing visual shapes are much different than those for haptic shapes, we might conclude that the neural underpinnings of these two abilities are substantially different (“late-convergence model”). Furthermore, mistakes in object recognition can be informative. If there is a distinct difference in the types of mistakes made when identifying shapes using the two different modalities, this would be evidence that the pathways responsible may be substantially different.

**Figure 1.**
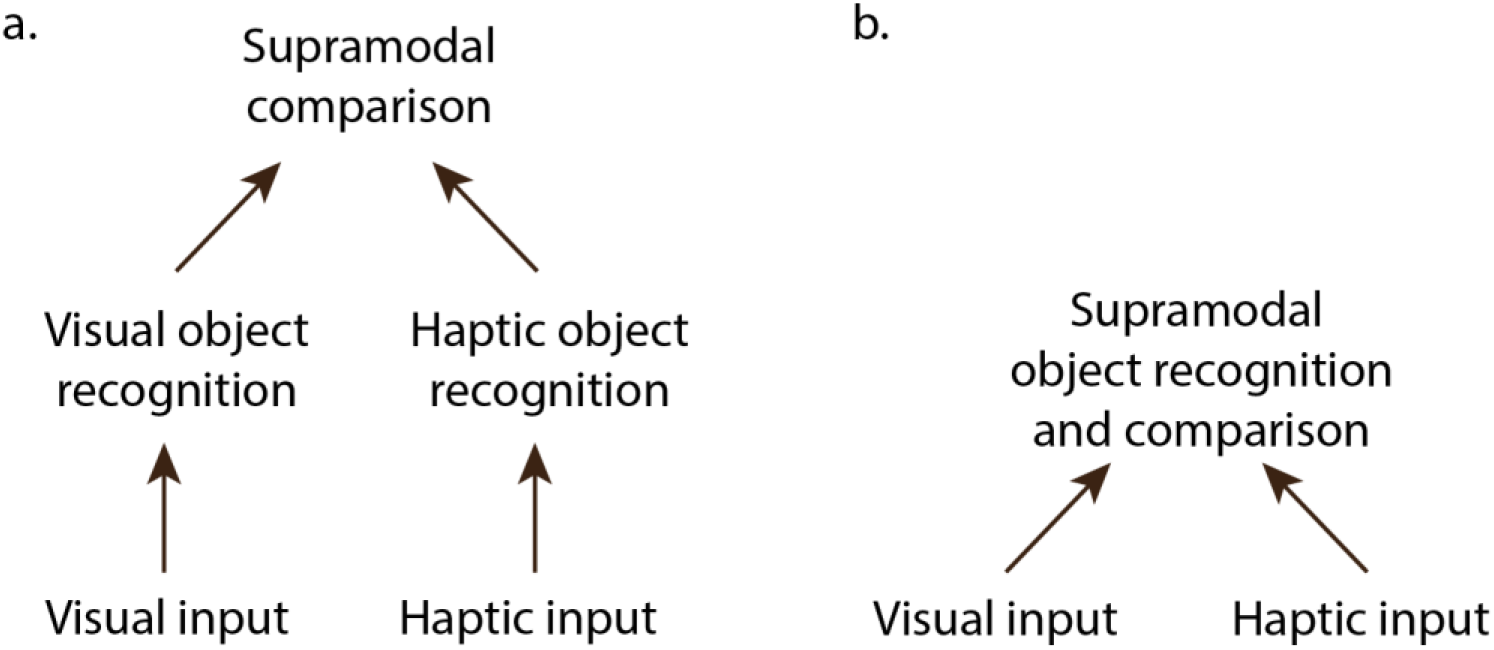
Schematized illustration of alternative hypotheses. **a**. Late-convergence model. One possibility is that shapes are recognized independently within unisensory processing areas and, if necessary, compared in some higher “supramodal” processing area. **b**. Early-convergence model. Alternatively, object recognition for the two modalities may use shared neural machinery following basic feature extraction (e.g., lines and curvature) in unisensory areas. (Adapted from Lacey et al., 2009)

We recruited human participants to perform a one-back shape matching task which involved determining whether a given two-dimensional abstract shape is the same or different as the shape which was presented seconds earlier. This was done both within modalities (i.e., comparing visual with visual, haptic with haptic) and across modalities (i.e., comparing visual and haptic shapes). This necessitated the design and production of a new device capable of quickly and reliably presenting physical objects from a large inventory (see Methods).

Our primary finding is that it is surprisingly simple to predict the subjective similarity of two abstract shapes for a given modality, but that similarity is modality specific. That is, two shapes which may appear visually similar may not necessarily feel haptically similar. Together, we take these results as evidence in support of a late-convergence model of cross-modal shape recognition.

## Methods

Human participants (n=10, 8 female) were recruited to perform a visual-haptic matching task lasting approximately one hour and were paid $15. Methods were approved by the Brown University Institutional Review Board. All participants had normal or corrected-to-normal vision. All participants were right-handed. One participant was excluded from this study due to miscommunication of instructions.

### Task

Participants performed a one-back task where they were asked to determine if the current stimulus was the same shape as the previous stimulus and report their answer by pressing the button corresponding to “same” or “different”. Stimuli were presented one at a time in blocks of 72 trials. Participants sat with their heads resting in a chin rest for all conditions, and all trials began with a fixation point appearing until fixation was acquired, then the fixation point disappeared and either a visual stimulus was presented or they were free to touch the haptic stimulus. Orientation of the visual or haptic shape was pseudorandomly chosen on each trial so that there was a 1:1:1 ratio between matching trials which were the same orientation, trials which were rotated 90 degrees left or right, and trials which were rotated 180 degrees. Each trial was pseudorandomly chosen as “same” or “different” so that each block of 72 trials used 48 unique shapes and had 24 “same” trials. Within those constraints, there was no limit on consecutive “same” or “different” trials. For example, it was possible (though exceedingly unlikely) to have the same shape presented 5 times in a row. Consistent with previous experiments comparing visual and haptic stimuli (Newell et al., 2001; Lacey et al., 2007, 2009; Tabrik et al., 2021), participants were given double the time to explore haptic (6 s) as visual (3 s), after which point the stimuli were removed. They could report their decision at any time during or after the stimulus presentation.

Each block was one of three types: visual-only, haptic-only, or alternating. “Alternating” blocks alternated between visual and haptic stimuli (Figure 2c). These three block types provided four conditions, based on the within- or cross-modal comparison being made: VV and HH correspond to the within-modal visual and haptic comparison trials, respectively. For VH trials, the visual shape presented on the previous trial is compared with the haptic shape on the current trial. For HV trials, the haptic shape presented on the previous trial is compared with the visual shape on the current trial. Each participant completed two visual-only blocks, two haptic-only blocks, and four alternating blocks, yielding 144 trials of each of the four conditions. The order of these eight blocks was counterbalanced across participants so as to avoid any order effect. Two-thirds of trials were “different” while the remaining 1/3 were “same”. Participants were given approximately 20 practice trials for each of the three block types before beginning data collection.

**Figure 2.**
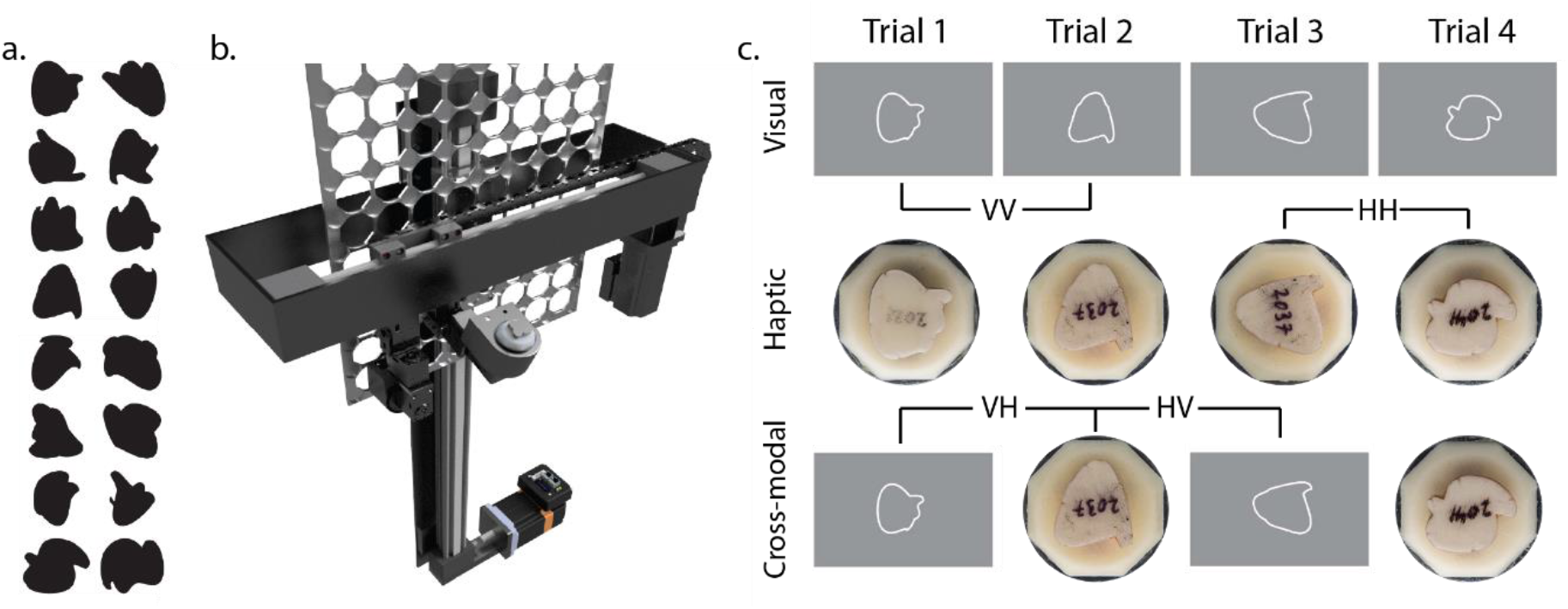
Stimuli and task. **a**. Randomly selected 16 of 48 shapes used in this task. **b**. Rendering of device used to present physical shapes to participants for haptic trials. **c**. Task conditions. Each participant performed the 1-back matching task in three block types: visual-only (top), haptic-only (middle), and cross-modal (bottom). These three block types yield four conditions: VV, HH, VH, and HV. Block types were completed in pseudo-random order, counterbalanced across participants. Each block consisted of 72 trials.

### Visual stimuli

On each visual trial, an abstract white-outlined shape, centered at fixation, was presented on a uniform grey background. Each shape was scaled such that it was the same size (~3 degrees visual angle) as the haptic representation of that same shape and rotated to one of four orientations, spaced in 90-degree increments. The shapes were constructed in a manner similar to Sigurdardottir & Sheinberg (2015). Two or three 2-dimensional “blobs” were generated using three randomly chosen control points connected by splines. Those blobs were then overlaid on each other and filled to produce a unique shape composed of multiple rounded edges (from the splines) and sharp corners (where 2 blobs intersect). Only unions of blobs which yielded a single filled shape were allowed. Examples of shapes used in the experiment are shown in Figure 2a. Stimuli were presented on an LCD monitor (Display++, Cambridge Research Systems) with 100 Hz refresh rate and eye position was tracked using an EyeLink eye tracker operating at 1 kHz to ensure fixation at the start of each trial.

### Haptic stimuli

A custom apparatus was designed and built (Figure 2b) to hold an inventory of up to 80 unique objects and present any one or two of those 80 objects at a given time (only one was presented at a time in the present experiment) to either the left or right hand. For this study, subjects explored the haptic shapes with their left hand, using their right hand to press one of two buttons indicating “same” or “different”. The haptic stimuli were positioned such that they could not be seen by the participant. The haptic presentation system includes an X-Y slide system (Igus drylin linear actuators, East Providence, RI, USA) driven by stepper motors (Applied Motion, Watsonville, CA, USA) used to position the inventory panel and two arms, each composed of three servo motors (Robotis Dynamixel, Seoul, South Korea) used to retrieve, present, and return objects.

Each object could be independently rotated in-plane for presentation at any angle. Each object was wrapped in 6 mm conductive foil tape (Adafruit, New York, NY, USA), and that tape was divided into six sections which were monitored at 100 Hz using a 12-channel capacitive touch sensor (six channels available for each of two objects, MPR121, Adafruit), enabling us to know when and where a shape was being touched by the participant.

On each haptic trial, a physical stimulus was presented to the left hand of participants at a comfortable position where their hand would naturally rest with elbows on their chair's armrest, rotated to one of four 90-degree positions. Haptic stimuli were simply an extruded version of the 2-dimensional visual stimulus. Each 2D shape was first scaled such that the maximum extent was 25 mm, then extruded 5 mm in depth using CAD software (Autodesk Fusion 360) and 3D printed (Mojo 3D printer, Stratasys, Eden Prairie, MN, USA). After 3D printing, the perimeter of the shape was wrapped in foil tape and divided into six sections as described above. Each of those six touchpads was wired to a custom circuit board embedded within each object to make those touchpads electronically accessible. The entire object was then painted with conformal coating (MG Chemicals, Surrey, B.C., Canada) to give a smooth, uniform feel and protect the copper touchpads.

For this experiment, we specifically chose to present (extruded) two-dimensional shapes, recognizing that these are only a subset of the kinds of objects encountered in the real world. For the visual-haptic comparisons under study, a significant advantage of using extruded 2D (as opposed to 3D) stimuli is that all relevant shape information is available to both modalities from a single view. With complex 3D shapes, a single view cannot reveal the entire shape (because you cannot see the back of an object) but a participants' fingers would have access to that shape information, leading to a fundamentally different opportunity to perceive the shape.

When presenting physical stimuli (as opposed to digital stimuli rendered on a computer screen), care must be taken to avoid any possibility that the participant might gain additional helpful information from the sights or sounds generated by the presentation mechanism. For example, in the present same-different task, it would be trivially simple to perform perfectly just by listening to whether the machine picks up a new object (“different” trials) or not (“same” trials). Multiple steps were taken to address such confounds. First, the presentation device was obscured from view, thus providing no helpful visual information. Second, after each trial, when an object was returned to the panel holding the inventory of objects, it was returned to a new location. This prevented a participant from being able to guess the identity of an object by listening to the x-y travel of the machine. Third, on every haptic trial, whether it was a “same” (the same haptic stimulus needs to be presented on successive trials) or “different” trial, an object was always dropped off and a different object was picked up. The only difference was that one robotic arm was used to drop off and pick up a new object on “different” trials and the second robotic arm was used to drop off and pick up a new sham object on “same” trials. The sham object was not actually presented to the participant, but there was no visual or auditory cue available to tell if the real or sham object was presented, and thus no cue predicting if the current stimulus was the same or different from the previous stimulus was present. Post-experiment questionnaires confirmed that participants had not found any strategy that was helpful in predicting the identity of haptic stimuli.

#### Behavioral measurements and analysis

##### Shape measurements

One of the major goals of this study was to determine whether participants were more likely to mistake two different shapes as being the same shape if the two shapes were similar. The question then becomes: how do you define “similar”? Here, we selected a variety of metrics with which we can quantitatively evaluate and compare shapes that are intended to cover a wide range of plausible methods of comparison. We acknowledge that although we have employed a wide range of these metrics, our collection does not constitute an exhaustive list of possible metrics.

###### Distribution of angles

Each shape was defined by roughly 700-800 (depending on the perimeter length of each shape) points spaced at 0.1 mm increments. To calculate the distribution of angles for a given shape, the local angle at each of those points was calculated over a specified span. For example, for a span of 101 points, the angle at point p is calculated as the angle formed by the vectors from p to p-50 and from p to p+50. The distribution of all angles making up a shape is simply the cumulative density function (CDF) composed of these angles.

###### Aspect ratio

For a given shape, the x span (xmax – xmin) and y span (ymax – ymin) were calculated. Then, the aspect ratio at that rotation is calculated as xspan/yspan. The shape is then rotated at 1 degree increments and the aspect ratio is computed again at each orientation. The aspect ratio for the shape is determined to be the largest of the 360 aspect ratios calculated for each shape.

###### Area / convex hull area / compactness

Area was calculated using the built-in MATLAB function “polyarea”. The convex hull was determined using the built-in MATLAB function “convhull” and then the convex hull area was calculated using the function “polyarea”. Compactness is defined here as the area divided by the convex hull area.

##### Shape comparisons

###### Distribution of angles

To determine the similarity of two shapes, A and B, using this method, we computed the sum squared error between the CDF of angles of shapes A and B.

###### Area / convex hull area / compactness

As with aspect ratio, we defined the similarity of two shapes in these measures to be the difference squared of the relevant measure.

###### Turning distance

This was calculated using the built-in MATLAB function “turningdist” based on (Arkin et al., 1991). Briefly, turning functions are calculated for each shape as the angle of the counter-clockwise tangent as a function of the length of each segment of a shape which is then normalized to a common length. This yields a complete representation of a shape that has the advantage of being invariant to size and x-y translation, but the disadvantage (for our application) of not being rotation invariant because the starting position for each turning function is arbitrary. Thus, the distance between turning functions is calculated for all starting positions of one shape and the minimum distance is taken as the turning distance between two shapes.

###### Intersection over union

This was calculated using the built-in MATLAB functions “intersect” and “union”, with the area of intersection then divided by the area of union. Possible values range from 0 to 1, with 1 representing perfect overlap between shapes. This ratio is computed for the pair of shapes both for the actual orientations as presented (“@actual”) as well as at the optimal orientation that maximizes the overlap (“@optimal”) to represent the mental rotation a participant may be doing to attempt to align two shapes.

###### Aspect ratio

The similarity of two shapes' aspect ratios is defined as the difference squared between the two shapes' individual aspect ratios (@optimal method). Difference in aspect ratio is also measured under the assumption that no attempt at mental rotation is made (@actual), calculated as the aspect ratio of a bounding box with the long axis oriented the same as that for the shape on the previous trial. For example, if the shape presented on the previous trial had its long axis oriented 30 degrees right of vertical, a participant may feel the current shape for a long axis that is around 30 degrees right of vertical.

###### Hausdorff distance

Shapes are first overlaid with aligned centerpoints. Next, for a given point on shape a, the nearest point on shape b was determined and the distance between these points was calculated. This was repeated for all points on each shape and the maximum of these minimum distances is the Hausdorff distance. Hausdorff distance is small for shapes which very nearly overlap and increase with larger deviations. Because this distance is not rotation invariant, it is calculated for both the actual orientations of the two shapes being compared (@actual) as well as the optimal orientation where Hausdorff distance is minimized (@optimal) to account for possible mental rotation a participant may do to align two shapes.

#### Metric evaluation

The various metrics were evaluated using a generalized linear model (GLM) with a binomial distribution to assess the relationship between the similarity of a pair of shapes (as determined by the metrics described above) with the response of a participant (“same” or “different”) on each trial. We used the Akaike information criterion (AIC) provided by the GLM as the dependent measure to evaluate a given metric. To facilitate comparisons within a condition (e.g., VH), a “random” metric was introduced which was simply a uniformly-distributed random number assigned for each trial. For a metric to be considered predictive, it should provide at least an additional 3 units of AIC beyond the random metric (Burnham & Anderson, 2004). This use of AIC for assessment was particularly helpful when comparing the performance of metrics by themselves with models composed of multiple metrics, as it accounts for the likelihood that adding predictor variables will tend to improve model performance (purely by chance) by penalizing for the additional factors. Here we chose the model with the fewest variables that was not improved by at least 3 units by adding an additional variable.

#### Monte Carlo simulation

In order to determine whether a given touchpad was touched more or less than would be expected by chance, and thus whether participants direct their haptic exploration towards particular features, we used a Monte Carlo simulation to form a baseline prediction of random touching. For each shape, 100,000 x-y points were randomly generated, each point representing a potential center point of a finger. If a given point was (a) outside the shape, (b) within 6 mm of an edge (representing the radius of a finger), and (c) not within 4 mm of an edge (representing the constraints on morphability of a finger when it touches a hard object), it was considered a touch. Any pad that was at least partially within 6 mm of the x-y point was considered “touched”. After 100,000 simulated touch points, the ratio of touches for each of the 6 touchpads was compared with the ratio of actual touches on those 6 touchpads to determine which pads were touched more or less than expected.

## Results

Ten participants participated in a one-back same-different task (Figure 2) during which they evaluated a series of abstract shapes to determine if each was the same as the last, regardless of changes in orientation. Each decision was signaled by pressing one of two buttons (“same” or “different”). Two-thirds of the trials were “different” trials and the remaining 1/3 were “same”. Each participant completed 144 trials for each of four conditions: VV (comparing two visual shapes), VH (comparing a haptic shape on the current trial with a visual shape on the previous trial), HV (comparing a visual shape on the current trial with a haptic shape on the previous trial), and HH (comparing two haptic shapes).

### Analysis of within-modal and cross-modal conditions

We used a multilevel linear model to analyze the repeated measures data. As is quite obvious from the results presented in Figure 3, the individual modalities used for the shape comparison (i.e. VV, HV, VH, VV) had a significant effect on percent correct performance (χ^2^ (3) = 55.84, p < 0.0001), sensitivity as measured using d-prime (χ^2^ (3) = 61.84, p < 0.0001), and reaction time (χ^2^(3) = 83.60, p < 0.0001).

**Figure 3.**
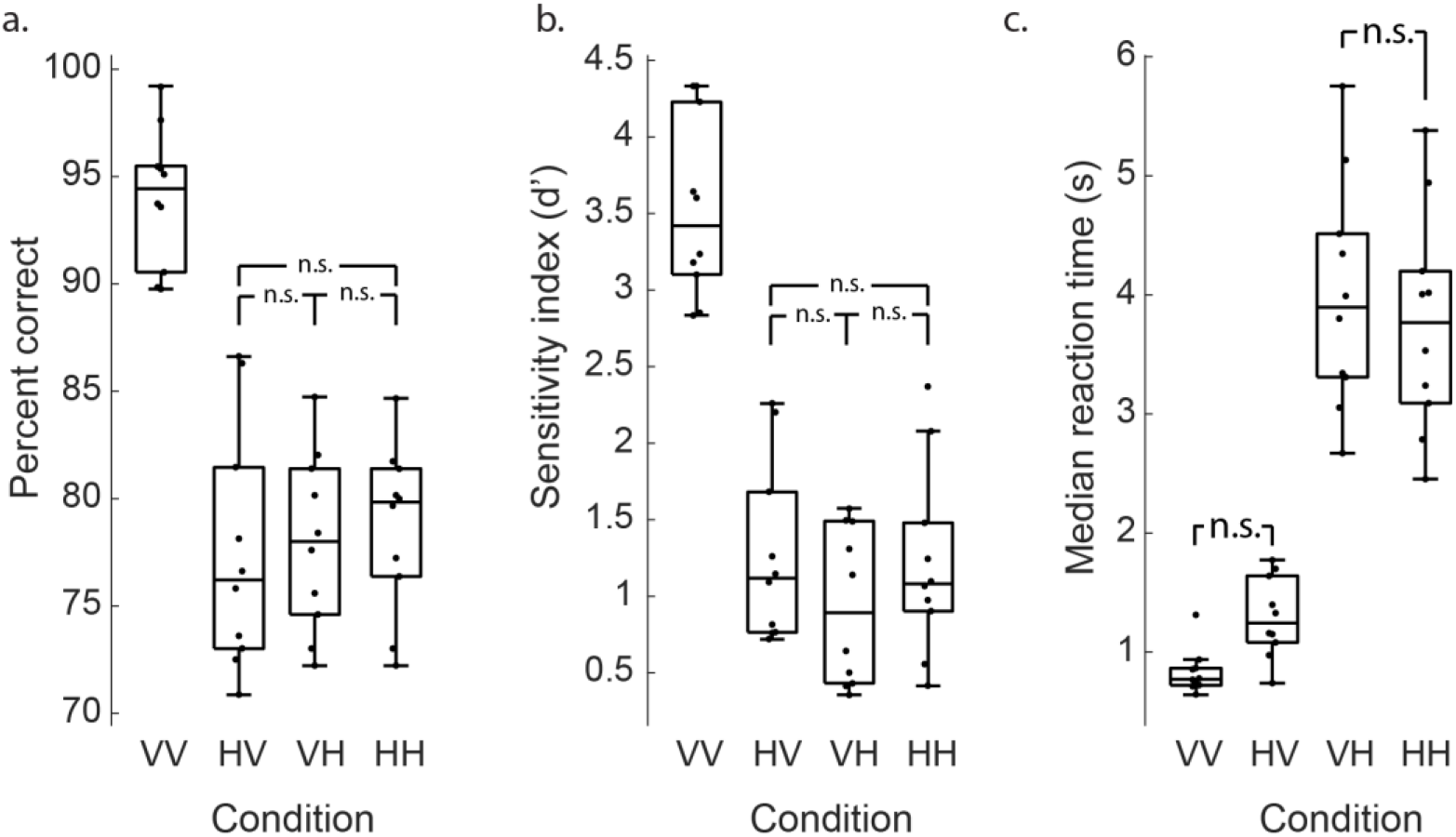
Task performance. All comparisons not labeled “n.s.” were significantly different (Tukey contrasts, p<0.005). **a**. Box plot showing performance (median, interquartile range, range) of all participants on each condition. Participants performed best on within-modal visual comparisons (VV). **b**. Performance using d-prime measure. **c**. Median reaction time (RT) for each participant for each condition. Cross-modal responses tended to be slower than within-modal comparisons (p=0.029), though each of the two cross-modal conditions (HV and VH) were not significantly slower than the comparable within-modal conditions (VV and HH, respectively).

For both the percent correct measure and the d-prime measure, orthogonal contrasts showed that the modality of the first stimulus in the comparison, the modality of the current stimulus in the comparison, and whether the comparison was cross- or within-modality all had a significant effect. In particular, if the prior stimulus was visual, accuracy and sensitivity were both significantly higher (for percent correct: b = 3.97, t(27) = 6.43, p < 0.0001; for d-prime: b = .496, t(27) = 5.63, p < 0.0001). Similarly, if the current stimulus was visual, accuracy and sensitivity were also significantly higher (for percent correct: b = 3.72, t(27) = 6.04, p < 0.0001; for d-prime: b = .662, t(27) = 7.52, p < 0.0001). Finally, performance in the within-modal, compared to the cross-modal, conditions was better (for percent correct: b = 4.30, t(27) = 6.98, p < 0.0001; for d-prime: b = .637, t(27) = 7.23, p < 0.0001). Using Tukey contrasts for multiple comparisons for performance and sensitivity, we also found that the VV condition was different from all three other conditions (all z-values < −9.0, p-values < 0.0001), but that none of the other conditions differed significantly from each other.

Contrasts for the reaction time data provide a slightly different picture. Not surprisingly, as we found for performance and sensitivity, the modality of the current stimulus significantly affected reaction times, with responses to a currently presented visual stimulus being significantly faster than a current haptic stimulus (b = −1407, t(27) = −18.82, p < 0.0001). However the modality of the prior stimulus did not have a significant impact on reaction times (b = −58.5, t(27) = −.78, p = 0.44). We did find, though, that the within-modal conditions were overall faster than the cross-modal conditions (b = − 172, t(27) = −2.31, p = 0.029). We again used Tukey contrasts to compare the four conditions to each other, and we found that the VV condition differed from the VH and HH conditions (z values < −14) but that although VV was slightly faster than the HV, the effect was only marginally significant (estimate = 462, z = 2.3, p = 0.097). When comparing VH to HH, the within-modal haptic condition was faster (estimate = 228), but the difference was not significant (z = −1.135, p = 0.66). We note that the reaction time cost for comparing HV vs VV is higher than comparing VH to HH. This suggests that translating a haptic representation in memory to compare to a visually presented match is more demanding than translating a visually stored shape into a haptic space.

### Effects of shape rotation

For match trials, the same shape was presented twice sequentially but the orientation of that shape could differ on each of the two presentations. This was primarily to curtail certain undesirable strategies (e.g., feeling only the top-left of each shape and comparing that small section between shapes) but it also allowed for an evaluation of the impact of rotation on recognition in different modalities. To assess the effects of rotation, and in particular to ask if rotation had a different impact on the recognition of shape as a function of modality, we modeled the percent correct and reaction time data using a factorial repeated-measures GLM (sensitivity could not be assessed as above as these were all match trials so we only have hits and misses). For this analysis, the first factor included levels for condition (VV, HV, VH, and HH). The second factor was stimulus rotation (0, 90, 180). To test for the overall effect of each factor we added them one at a time to the baseline model followed by inclusion of their interaction.

Results for percent correct performance are shown in Figure 4. Considering hit rate, the addition of each factor and the interaction significantly improved the model fit (modality: χ^2^(3) = 28,4, p < 0.001; rotation: χ^2^ (2) = 13.9, p = 0.0009; interaction: χ^2^ (6) = 30, p < 0.0001) and reaction time. This implies that performance across the four conditions differed, and that performance depended on the orientation of the sample and match stimulus. The significant interaction indicates that the effect of rotation itself depends on the modality.

**Figure 4.**
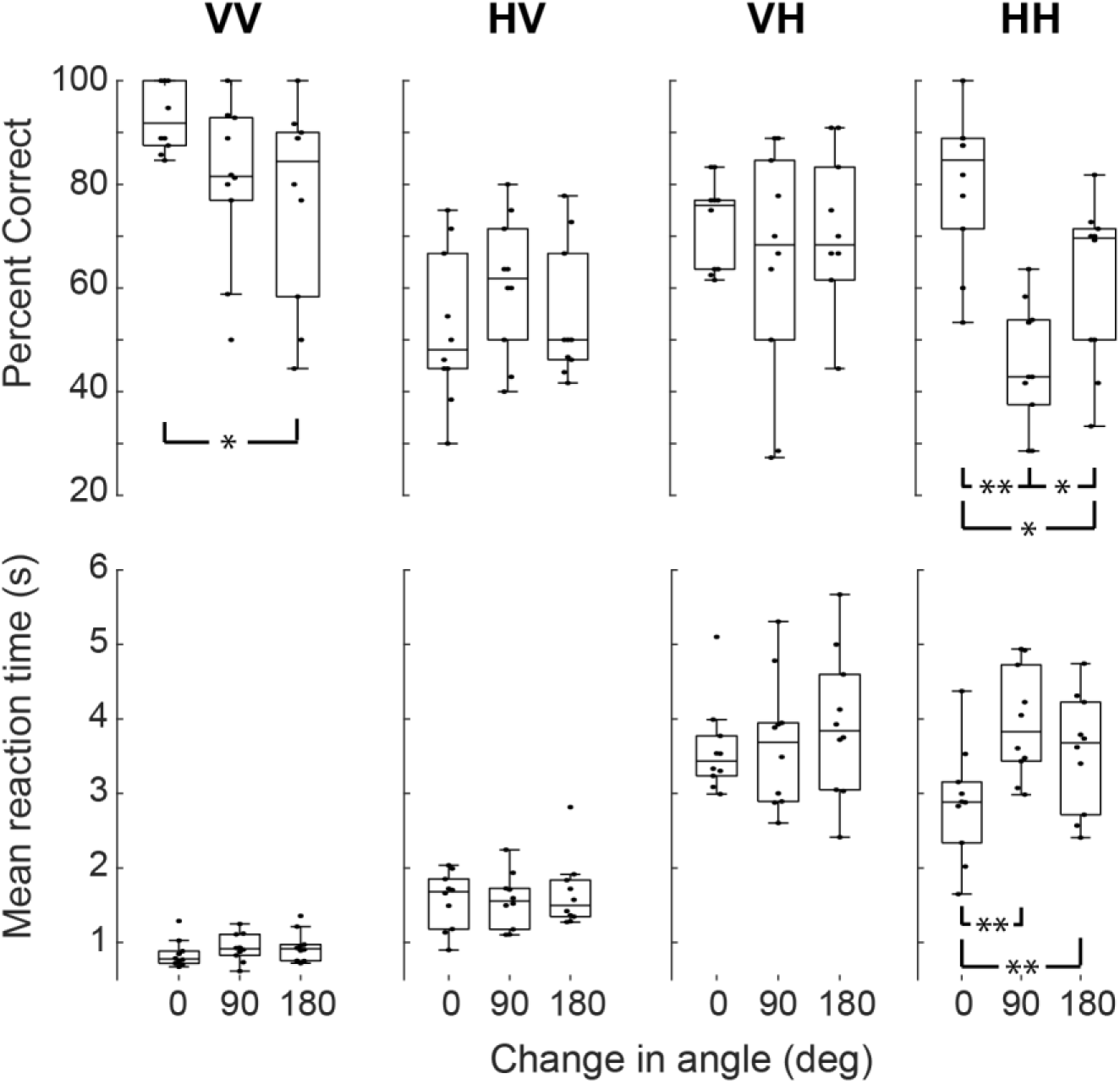
Performance with rotation of match shapes. Within-modal rotation tends to lead to more mistakes and longer RTs, while no such penalty is seen for cross-modal comparisons. (* p<0.05, ** p<0.005, all others p>0.05)

Specific contrasts help us better understand these trends. For the modality factor, we included orthogonal contrasts for “visual current” (VV & HV vs VH & HH), “cross modal” (VV & HH vs HV & VH) and “unimodal pathway” (VV vs HH, ignoring the cross-modal conditions). For the rotation factor, we included contrasts for “rotated” (0 vs 90 & 180) and for “degree of rotation” (90 vs 180, ignoring 0). All of these factors, except the last, were significant (visual current: b = −.06, t(27) = 2.72, p = 0.011; cross-modal: b = −0.05, t(27) = 3.45, p < 0.005; within-modal pathway: b = 0.34, t(27) = 5.34, p < 0.0001). These significant contrasts support the idea that regardless of rotation difference, performance on match trials is better when the current stimulus is visual, when the comparison is made within modality compared to across modality, and, for unimodal trials (VV and HH) that visual comparisons are more accurate than haptic comparisons.

For the rotation contrasts, we found a significant effect of rotation away from 0 (rotated: b = − .11, t(27) = 4.41, p < 0.0001) but no overall difference between the 90 and 180 degree rotation (degree of rotation: b = −0.02, t(27) = .75, p = 0.46). This supports a model where shape comparisons are orientation sensitive, as one might expect, with rotation between sample and match reducing performance. However, we also found a significant interaction between the cross-modal contrast and rotation away from 0 on performance (b = 0.1, t(27) = 4.14, p < 0.001). This suggests that orientation differences between sample and match have greater impact on performance for within-modal comparisons compared to cross-modal comparisons. Said another way, cross-modal recognition performance appears more invariant to rotation, a finding that accords with previous studies of visuo-haptic recognition (Bülthoff & Edelman, 1992; Newell et al., 2001; Lacey et al., 2007, 2009; Andresen et al., 2009).

Finally, we analyzed the effect of rotation specifically within the unimodal VV and HH conditions. Looking at Figure 4, it appears that the effects of rotation were more pronounced for the within-modal haptic condition compared to the within-modal visual condition, and that the patterns of the effect were distinct. Using general linear hypothesis testing we compared the 0-90, 0-180, and 90-180 rotation conditions within the VV and HH conditions. This analysis revealed that the effects of rotation for the visual matching trials were incremental and monotonic, (0 vs 90: b = .13, z = 2.149, p = 0.06; 90 vs 180: b = .03, z = .59, p = 0.55; 0 vs 180: b = 0.16, z = 2.74, p = 0.025). This suggests that for visual matching, performance degraded with increasing rotation. Rotation affected the haptic condition in a very different way. Rotation away from the original orientation by 90 degrees dramatically affected performance (0 vs 90: b = .35, z = 5.88, p < 0.0001) but a further 90-degree rotation actually improved performance (90 vs 180: b = −.16, z = −2.70, p = 0.025). We return to this distinction below, as we consider which specific shape dimensions appear critical for visual and haptic recognition and how these may differ. Consistent with the match performance measures, each factor and their interaction significantly improved the model fit for reaction times (modality: χ^2^(3) = 86.9, p < 0.001; rotation: χ^2^(2) = 16.3, p = 0.0003; interaction: χ^2^(6) = 38, p < 0.0001). We can infer that modality, rotation, and their interaction are all significant predictors of reaction times.

Results of the contrast analyses revealed that reaction times are, as is evident from Figure 4, significantly faster for trials where the current stimulus is presented visually (visual current: b = −1068, t(27) = 12.77, p < 0.0001). Reaction times for cross modal trials were also significantly slower than unimodal trials (cross modal: b = 241, t(27) = 4.08, p < 0.0001). Rotation of a stimulus between sample and match also significantly slowed responses (b = 306, t(27) = 5.12, p < 0.0001). Interactions between the modality and rotation factors also proved significant. The interaction between cross-modality and rotation was highly significant (b = −180, t(27) = 3.01, p < 0.005), which corroborates the performance results presented above, again suggesting that effects of rotation are less pronounced for cross-modal comparisons compared to within-modal comparisons. A significant interaction between the cross-modal contrast and the degree of rotation (90 vs 180) conditions (b = −182, t(72) = 2.62, p = 0.01) indicates that the 90 degree rotation slows responses for the within-modal condition but not the cross-modal condition. We also observed a significant interaction between the type of within-modal trial (visual/VV or haptic/HH) and any rotation (b = −620, t(27) = 2.59, p = 0.012) as well as the degree of rotation (b = − 571, t(72) = 2.06, p < 0.05). We note that although these interactions suggest that the visual and haptic within-modal comparisons are differentially affected by stimulus rotation, their interpretation is made difficult by the large difference in overall reaction time between the VV and HH conditions (see Discussion).

### Predicting behavior based on shape metrics

The behavioral differences observed on the same task in different modalities led us to model the behavior to better understand the critical features of shapes participants used to complete this task and to compare these critical features for both visual and haptic shapes. To this end, we chose eight “shape metrics” (Figure 5) which provide a variety of methods for quantifying the similarity between two shapes. Based on these shape metrics, we could then predict if two shapes were likely to be conflated. By comparing the success of these various shape metrics in predicting behavior in each modality condition, we can gain insight into how shapes are evaluated by vision and touch. Details of each of these shape metrics are provided in the methods section.

**Figure 5.**
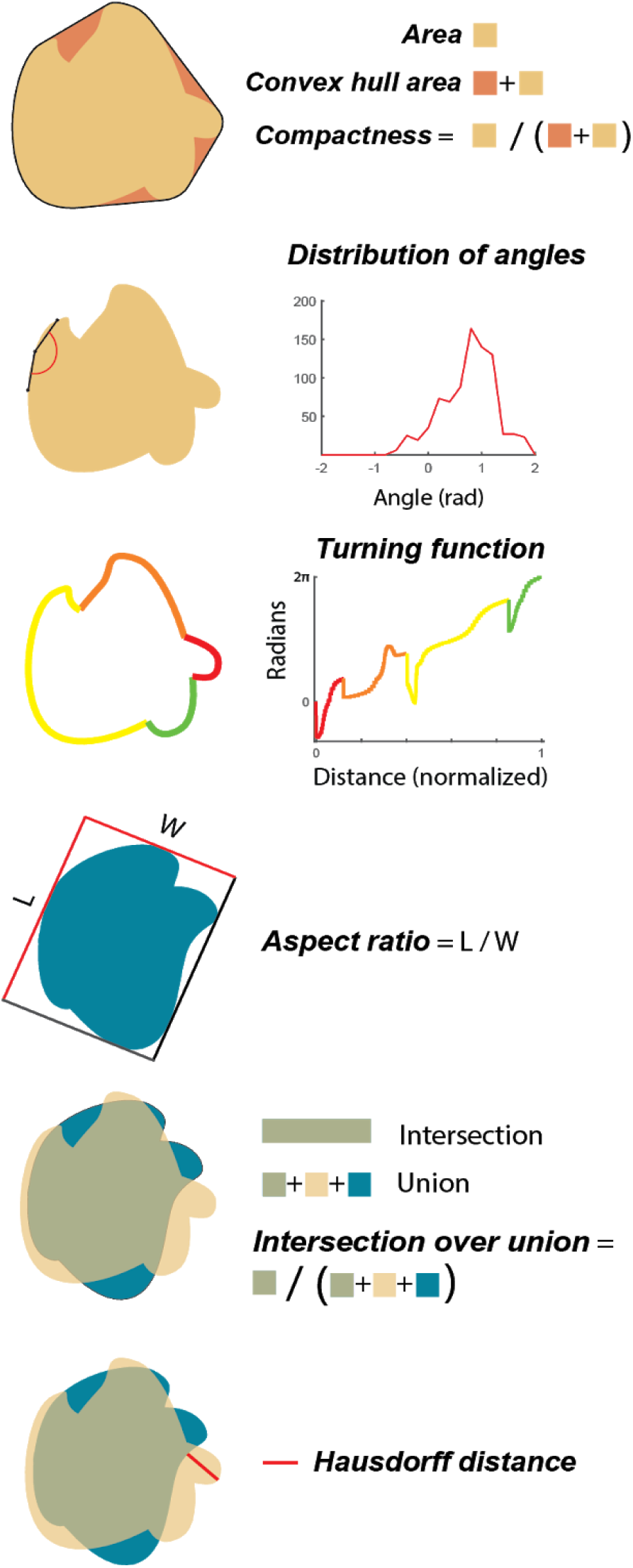
Shape comparison metrics. The metric “area” is the area encompassed by a shape. Convex hull area is the area encompassed by a boundary enclosing the shape with only convex turns. Compactness is the quotient of the area to the convex hull area. Distribution of angles represents all the angles which can be said to compose a given shape given a certain spread over which those angles are calculated. Turning function is a complete representation of a given shape using the tangent and length of each line segment composing that shape. This reformatting simplifies the comparison of shapes by being size and translation invariant. For explanatory purposes, an example shape is shown with color-coded sections. Aspect ratio (@optimal) is the largest ratio of length to width of a rectangle enclosing a shape. Intersection over union (@optimal) is the optimal overlap which can be achieved by overlaying one shape with another. Hausdorff distance (@optimal) is the maximum of all minimum distances between all points on one shape and all points on another shape, optimized by rotating one shape relative to the other to find the smallest possible Hausdorff distance for a given pair of shapes.

Figure 6 shows the performance of each model in fitting behavior for each of the four conditions. For within-modal visual comparisons, the single metric which best predicts behavior is distribution of curvature. This metric simply catalogs the various angles which compose a given shape without regard for the spatial relationships between those angles. Similarly, for VH trials (comparing a haptic shape on the current trial with the visual shape seen on the previous trial), distribution of curvature is once again the best metric. For HV trials, Hausdorff distance and intersection over union (@optimal) were most informative. Hausdorff distance is a simple and extremely sparse description of the differences between shapes, representing the maximum of all minimum distances between points on a pair of shapes. If two shapes can be oriented such that their boundaries nearly overlap, Hausdorff distance is small. However, if one of the two shapes has a large protrusion, but the shapes are otherwise identical, Hausdorff distance is large. Intersection over union is the ratio of overlapping to non-overlapping area shared between two shapes. Finally, behavior on within-modal haptic trials is best described by the compactness metric. We can think of this metric as a measure of the area of a shape's concavities (e.g., a circle is very compact because there are no concavities, whereas a starfish shape would not be compact). Considering the physical limitations inherent to manual evaluation of an object, it is not so surprising that concavities are particularly salient in haptic exploration.

**Figure 6.**
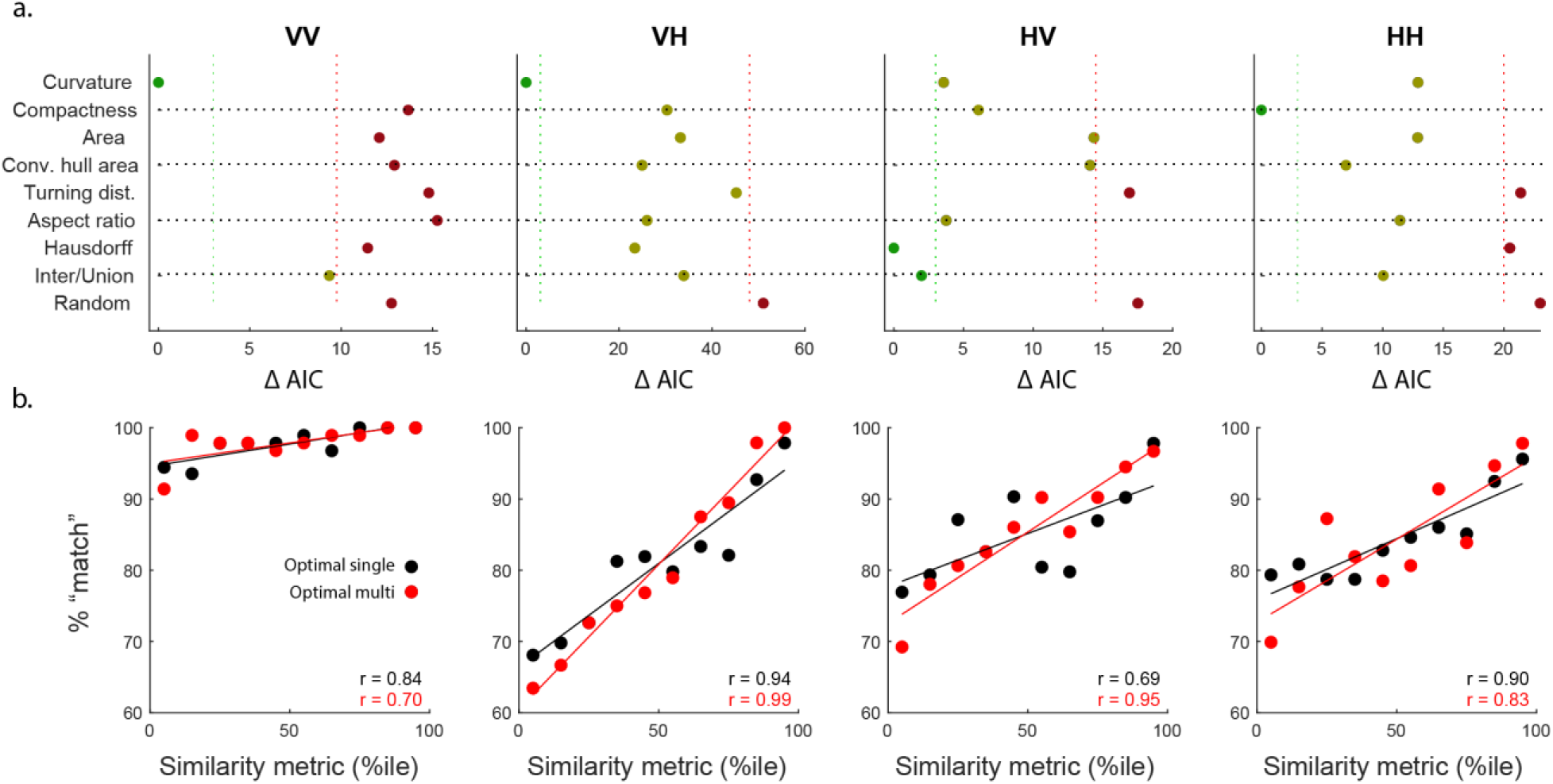
**a.** Performance for each similarity metric for each condition. Horizontal axis is the difference between AIC of each individual metric and the best metric for that condition. Vertical dashed green lines indicate the threshold for what can be considered no different from the best model, while vertical dashed red lines indicate the boundary for metrics which can be considered no different from chance. **b.** Behavioral performance on non-match trials correlates with the similarity of those shapes according to the most successful metric for that condition (Optimal single, black). As the difference between two shapes increases, the probability that a participant will correctly label them as “non-match” increases. In all four conditions, the use of additional factors significantly improved predictive performance although not necessarily leading to an increase in the correlation seen here. The optimal multi-metric for VV and HH used two factors, while the optimal multi-metric for HV and VH used three factors.

For a clearer sense of how well these metrics predict behavior, Figure 6b shows the relationship between the shape difference (according to the best metric for that condition) and the performance of participants. It is important to note here that the best metric was not chosen according to which metric produces the strongest correlation, but rather according to which best predicts the choice of the participant on a trial-by-trial basis. Nevertheless, there is a clear, nearly monotonic, relationship in each condition in the expected direction: shapes that are more dissimilar according to a similarity metric are more likely to be labeled “different” by a participant (Pearson's r, VV: r=0.84, p=0.002; VH: r=0.94, p=0.0001; HV: r=0.69, p=0.026; HH: r=0.90, p=0.0004). Although the trend is less pronounced in the VV condition, likely because of the near ceiling performance, the correlation is still highly significant. In all cases, performance for the most dissimilar shapes is nearly perfect. Figure 6b also shows the performance of the metrics when combined (red). Using Akaike Information Criteria (AIC) to “punish” models with added complexity, we found that the optimal models for each of the two within-modal conditions were best fit by combinations of two metrics, while each of the cross-modal conditions were best fit using combinations of three metrics. In all cases, these multi-metrics were better able to predict behavior even after accounting for the added factors, though this does not necessarily result in a stronger correlation coefficient. For VV trials, the best multi-metric was a combination of curvature and intersection over union while for HH, it was compactness and hull area. Interestingly, for both VH and HV, the exact same 3-factor multi-metric was best: the combination of aspect ratio, curvature, and Hausdorff distance.

Three of the metrics described here were each calculated using two different methods, described here as @optimal and @actual. The intuition here is that we do not know *a priori* if behavior in this one-back matching task is better modeled by assuming participants are performing mental rotation (as they should, they were instructed to ignore rotation) or not. Ideally, participants would have a perfect recall of the shape presented on the previous trial and would have the ability to compare that shape with the shape presented on the current trial at all possible orientations and evaluate the similarity at each of those orientations. If there is any possible orientation where the shapes are a match, then the response is “same”, otherwise “different”. However, we know that mental rotation abilities are imperfect (Shepard & Metzler, 1971; Gauthier et al., 2002) (Figure 4) so it may be that performance is highly dependent on the exact orientation at which those shapes happen to be presented. If they happen to be oriented, for example, such that they both have a protrusion on top, they may be labeled “same”, while if those same shapes are presented with protrusions on opposite sides they may be labeled “different”. Metrics which assess the similarity between two shapes at the optimal alignment which maximizes their similarity are labeled @optimal, while metrics which assess similarity at the actual orientations in which they were presented are labeled @actual.

Interestingly, we found a clear difference between within-modal and cross-modal conditions in terms of whether they were better fit by @actual or @optimal metrics (Figure 7). Within-modal behavior was better described by @actual metrics and cross-modal behavior was better described by @optimal metrics. This implies that mental rotation is costly or difficult when comparing shapes within the same modality but simple or even automatic when comparing shapes across modality. This provides independent confirmation supporting the results presented above (Figure 4) and reported previously (Lacey et al., 2009).

**Figure 7.**
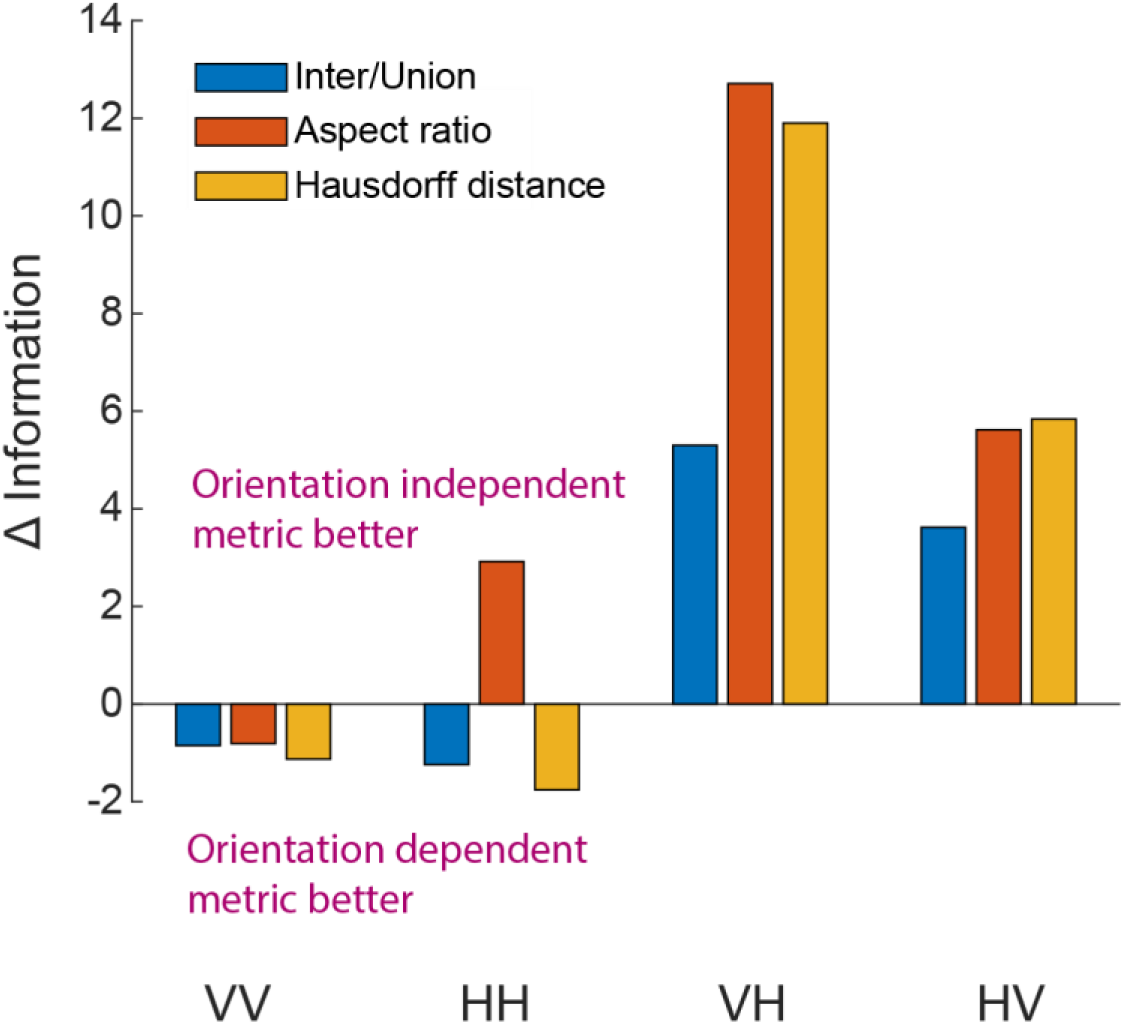
Within- vs cross-modal differences in behavior. Change in AIC between @optimal and @actual metrics show that within-modal behavior (conditions VV and HH) are better described by metrics which do not assume mental rotation (i.e., @actual metrics) while cross-modal behavior (conditions VH and HV) are better described by metrics which do assume mental rotation (i.e., @optimal metrics).

### Characterization of active exploration

We also sought to determine whether the way participants touch shapes differs based on experimental condition (Figure 8). Using the six touch sensors embedded in each shape, we quantified “dwell time” (duration of each touch of a touchpad indicating how quickly the hand moves around the object), “unique pads touched” (how many of the six pads are touched on a given trial), “simultaneous pads touched” (how many pads are touched at any given moment, corresponding to the number of fingers being used), and “total pad touches” (number of the six pads which were touched in the trial, an estimate of how much effort is spent on exploring a shape). No differences were found in dwell time, unique pads touched, or simultaneous pads touched between the HH and VH conditions, suggesting the basic strategy of how a shape is explored does not depend on the modality with which it is being compared. The only difference was found in the number of total pad touches, with more pad visits found in the VH condition (mean pad visits per trial HH: 12.1, VH: 12.5. K-S test, p=0.009). Interestingly, this difference in total pad touches between VH and HH resulted almost entirely from the specific condition where the same haptic shape was presented consecutively (i.e., “match” trial) at the same orientation, corresponding to the relatively quick reaction times in this condition (Figure 4, “HH” column, 0 change in angle). We conclude that people explore haptic objects the same whether comparing it to a previously presented haptic or visual shape, but that, as this comparison is more challenging, more time is spent carrying out that exploration.

**Figure 8.**
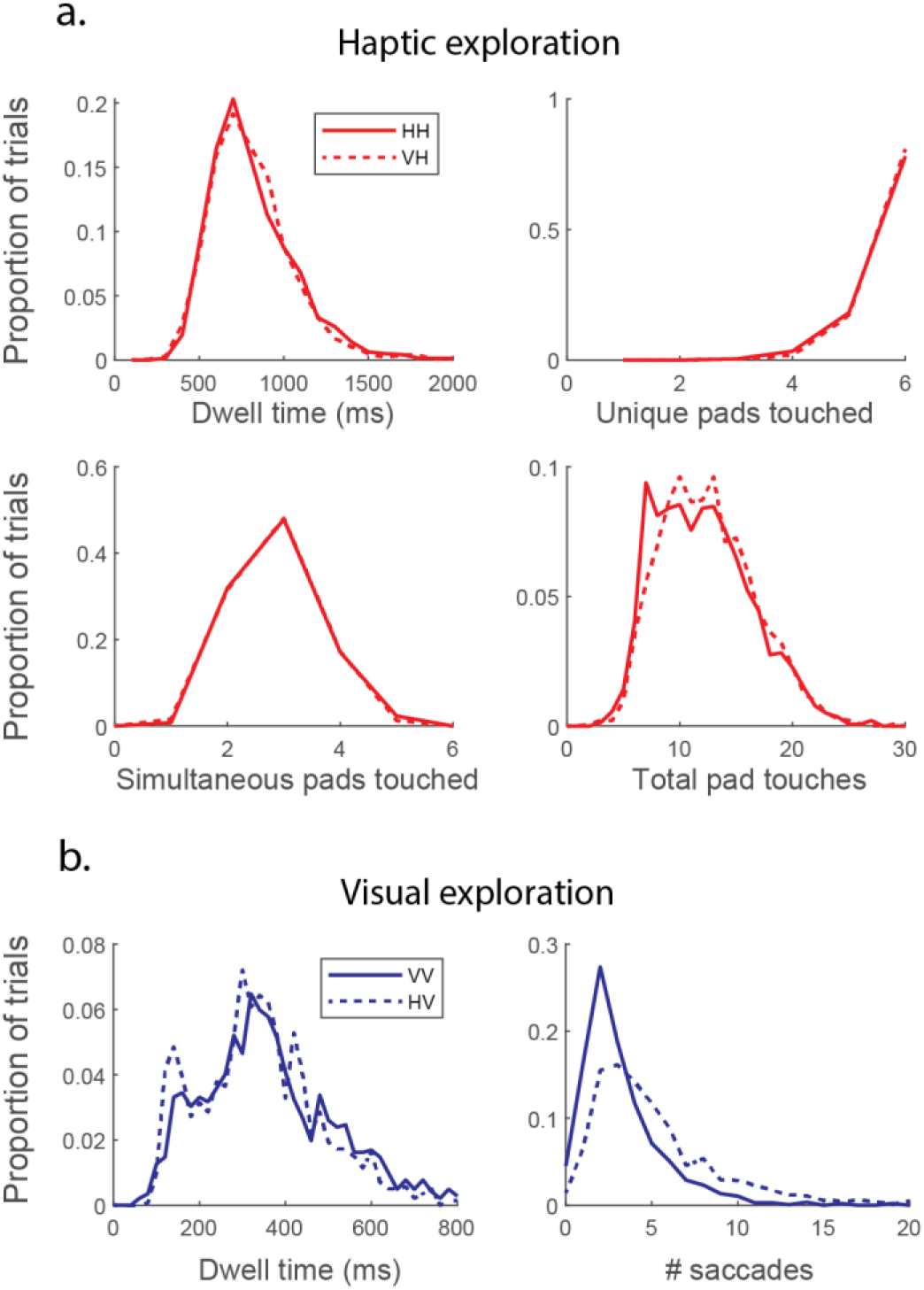
Statistics of haptic and visual exploration across all trials and participants. **a.** Haptic exploration of shapes is remarkably consistent regardless of condition. Participants spent the same amount of time touching each portion of the shape (“dwell time”), touched the same proportion of the shape (“unique pads touched”), and touched the same proportion of the shape at once (“simultaneous pads touched”). The only difference is found in the duration of exploration, leading to more touches in the VH condition (“total pad touches”, K-S test, p=0.009). **b.** Similarly with visual exploration, the biggest difference between conditions is that participants spend more time looking around the shape in the HV condition (K-S test, p=8e-47). Interestingly, the dwell time at each saccade endpoint is significantly shorter in the HV condition than in VV (K-S test, p=1e-5).

To compare this with the visual behavior, we evaluated dwell time (i.e., inter-saccade interval) and number of saccades per trial. Similar to the results for haptic exploration, the biggest differences were seen in the amount of time exploring (mean saccades per trial VV: 3.22, HV: 5.27; K-S test, p=8e-47), again corresponding to differences in reaction time (Figure 4). However, there was also a significant difference in the dwell time at each saccadic location with participants making more frequent saccades in the HV condition (mean dwell time VV: 405 ms, HV: 366 ms; K-S test, p=1e-5).

Finally, we asked if participants employ targeted exploration to focus on areas of a shape with more curvature or whether exploration appears more uniformly distributed (Figure 9). We used a Monte Carlo approach (details in methods) to estimate the predicted touches of each touchpad if haptic exploration were random and then compared that to actual touches of those touchpads (we would not expect an equal number of touches for each of the six touch pads because not every pad is the same length nor equally accessible to a finger). We found that, in both HH and VH conditions, the location of touches was not random but instead there was a significant relationship between the amount of time participants inspect a given touchpad and the amount of curvature in that area (Pearson's correlation. HH: r=0.32, p=4e-8; VH: r=0.28, p=1e-6), suggesting that participants intentionally focus on exploring complex and presumably “diagnostic” features.

**Figure 9.**
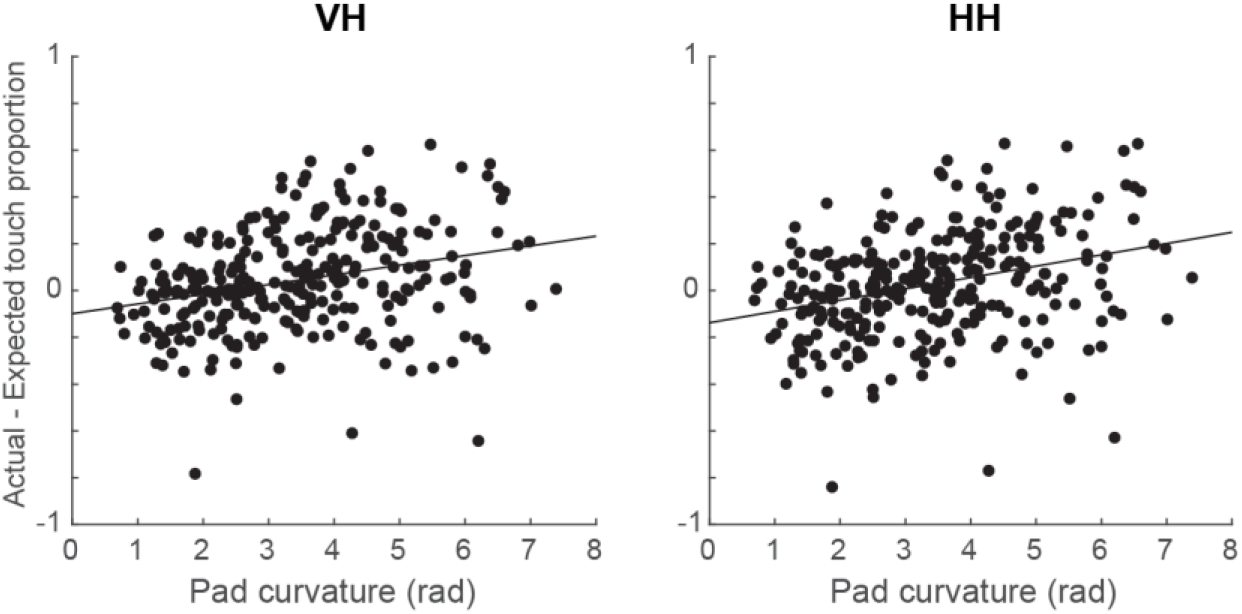
Regions with more curvature are touched more than expected based on Monte Carlo simulation. In both VH trials (left) and HH trials (right), touchpads with more curvature are touched more than would be expected by chance, whereas touchpads covering straighter portions of a shape are touched less than would be expected by chance. This indicates participants specifically target areas of high curvature for manual exploration. Each point represents one touchpad on one shape. Line is linear best fit. (Pearson's correlation. HH: r=-0.32, p=4e-8; VH: r=0.28, p=1e-6)

## Discussion

In this study, human participants performed a one-back shape comparison task where they were presented a continuous stream of shapes and asked to report if the currently presented shape was the same as the previously presented shape. Shapes were presented either on a computer monitor (visual) or by a robotic arm to the participants' left hand (haptic). Though our specific paradigm and the manner of presentation of physical objects were novel, we did confirm previous results found in visual-haptic research (Bülthoff & Edelman, 1992; Newell et al., 2001; Lacey et al., 2007, 2009; Andresen et al., 2009), showing that orientation is important when comparing shapes within a modality but not when comparing shapes across modalities. This increases our confidence that the results shown here are broadly applicable for visual-haptic research using different stimuli and different presentation methods.

The primary new findings in the present study are that (1) performance in this shape-matching task is predictable based on simple metrics which quantify the similarity between shapes, and (2) that the metrics which best predict behavior depend on the presentation modality. For example, the best metric for predicting a within-modal visual comparison is very different from the best metric for predicting behavior on a within-modal haptic comparison. This does not appear to result from simple differences in spatial acuity between the senses but rather from fundamental differences in the way shapes are processed.

These results may seem to contradict a recent study by Tabrik and colleagues (2021). In that study, a great deal of similarity was found between the chosen shape metrics and the self-reported measures of perceptual similarity in both within-modal visual and within-modal haptic comparisons (different groups of participants were used for these tasks, so no cross-modal comparisons were possible). A number of differences between that study and the present study could explain the discrepancy. First, it may be that self-report of perceived shape similarity on a 7-point scale is different from the “revealed perceptual similarity” obtained in the present study, where we analyze which shapes are confused for other shapes. For example, a participant may assess that two shapes are very similar to each other overall and yet the small difference could be quite salient such that they would never be confused for each other. Second, the Tabrik et al. study used shapes which were generated by evolving eight shapes from each of two related initial shapes using digital embryo algorithms. Because the shapes were all part of the same family, that may have encouraged evaluation of shape differences on a given set of dimensions which best describe the differences in that specific shape family but wouldn't necessarily describe differences in independently generated shapes. Third, the Tabrik et al. study allowed participants to manipulate the shapes (physically in the haptic condition, virtually in the visual condition) whereas the present study did not. This raises the interesting possibility that active manipulation of a shape may alter perceived similarity between shapes.

It is important to note that the metrics used here to quantify differences between shapes can be used predictively. That is, it should be possible to intentionally create shape sets which are difficult to differentiate visually or haptically. Furthermore, it should be possible to create stimulus sets which are difficult to differentiate haptically but easy to differentiate visually, and vice versa. Some previous work in this area has used post hoc analyses of behavior to group shapes by similarity, but does not provide a means of directly predicting perceived similarity in the absence of behavioral results (Huang, 2020). We sought here to develop models which are more readily interpretable and can thereby provide greater intuition (Rudin et al., 2021).

It is also important to emphasize that object familiarity may play a role in the extent to which different brain areas are involved in visual and haptic object recognition. Previous work (Deshpande et al., 2010; Lacey et al., 2010) has indicated that the networks involved in haptic object recognition are similar to visual object recognition only when the shapes are familiar. This may also explain the results found in Tabrik et al. (2021) where shapes were likely more familiar and greater similarity was found between visual and haptic processing as compared to the present study. Further work will seek to determine the extent to which stimulus familiarity impacts which metrics best predict human behavior.

Finally, this work suggests something about shape processing in the brain, more generally. Our initial hypothesis was that, to the extent that haptic object recognition recruits visual cortical areas for processing object shape, the same properties that are important for differentiating visual shapes should be important for differentiating haptic shapes. For example, visual shape recognition is thought to rely on combining activity of neurons in visual cortex sensitive to local curvature (Riesenhuber & Poggio, 1999; Serre et al., 2005; Yau et al., 2009; Pasupathy et al., 2020). This would predict that two different shapes with similar local curvature would be easily confused. If haptic shape recognition uses the same pathways, we would expect local curvature similarities to also contribute to mistakes of haptic shape recognition, particularly if we allow for different definitions of “local” based on the lower acuity of haptic vs. visual perceptions (i.e., measuring local curvature over various distances which should be optimized based on fingertip size). This does not appear to be the case. Rather, the metrics that work well for predicting haptic-haptic shape comparison appear fundamentally different from those that work well for predicting visual-visual shape comparison. This suggests that pathways for within-modal haptic shape processing may exist somewhat independent of visual processing. While these pathways have not yet been fully mapped, the device developed in this study for presenting haptic objects could be used to explore these circuits more systematically than has been possible in previous studies.

The finding that behavior in the cross-modal conditions is best fit by models which combine three metrics, while behavior in the within-modal conditions is best fit by models which only combine two metrics, further bolster the view that processing across modalities is fundamentally different than processing within modality. The observed increase in complexity required to explain behavior may reflect an increase in complexity of the networks involved and the need for interactions between these, which can perhaps be short-circuited for within-modal comparisons, most notably in the absence of rotation. Further work, particularly using electrophysiology and neuroimaging techniques, should prove useful in elucidating the areas which are involved in these varying shape recognition scenarios.

